# FoodEstNet: Estimating True Food Consumption with Machine Learning

**DOI:** 10.1101/250506

**Authors:** Darlington A. Akogo, Joseph B Danquah

## Abstract

We developed a Machine Learning/Artificial Intelligence model that estimates how much of a food type a person truly consumes. People tend to underestimate how much they consume which makes the work of nutritionists and dietitians difficult since they rely on food estimates for food portion size control and nutritional management of diseases. We trained an XGBoost model to estimate how much a patient truly consumes based on Age, Sex, BMI, socioeconomic status and perceived consumption.

## Introduction

The researchers, Livingstone and Pourshahidi (2014) reported that consuming more than the needed portion size at meal times leads to over consumption of calories and energy, especially from energy dense foods. Consumption of large portion sizes may be a risk for obesity; an epidemic that has been linked to the development of non-communicable diseases, (NCDs) (Cecchini et al, 2010; Schwartz & Byrd-Bredbenner 2006). The increase of obesity prevalence has been partly attributed to the increased consumption of food portion size which is largely due to overestimation. Insufficient consumption of food portions has been linked to undernutrition and undesirable health consequences especially in pregnant women and children up to two years (Black et al., 2008; Benson and Shekar, 2006). Intrauterine growth retardation as a result of maternal undernutrition results in infants with low birth weight (LBW). Low-birthweight infants and poorly nourished children are at risk of developing NCDs such as cardiovascular diseases, hypertension and diabetes in their adult life (Barouki et al, 2012 and Delisle, 2002). Nutrition scientists claim that underestimation of true calorie intake by people ranges from 10% to 40%, with a typical underestimation of 30% (Marion & Maiden, 2014). The underestimation according to the researchers are linked to factors such as age, sex, body composition, and socioeconomic status (Bowman, Lino, Gerrior, & Basiotis, 1998). Such inaccuracy to estimate actual food portion consumed is a major barrier to the achievement of food portion size control and its effect in the nutritional management of diseases. In our research, we present FoodEstNet, an XGBoost model that takes in 11 demographic, socioeconomic status attributes and perceived consumption as input and then outputs an estimation of actual consumption. FoodEstNet is trained and tested on a dataset with 4,800 data samples and 11 features/attributes with one output. Included in the input and dataset is the attribute “Food Type” which includes commonly consumed carbohydrate foods in Ghana (et al Boateng, 2014), Chocolate drink, white bread, boiled rice, Ga kenkey, granulated sugar, corn porridge, waakye, gari, boiled yam, boiled green plantain, banku and fufu. So, for each input run, attributes pertaining to demographics and socioeconomic status are inputted, a specific food type among the 12 food types is chosen, perceived consumption amount is finally inputted also and FoodEstNet tries to estimate the actual consumption amount. After training, we tested the model separately on the test data split using Root Mean Square Error and Explained Variance Score functions as metrics.

### Portion size and the obesity epidemic

Globally, consumption of large portion sizes of energy dense foods such as fats, refinedcarbohydrates, and low fiber foods are believed to be on the increase (WHO, 2012a; Vorster, etal, 2011). These unhealthy dietary practices as well as decreased physical activity have been linked to the development of overweight and obesity (Popkin et al, 2012). Positive differences in energy intake and output as the cause of the obesity epidemic (Stubbs and Lee, 2004). The relationship between increased portion sizes, overweight, obesity (Wansink, 2011; Wansink, 2008; Schwartz and Byrd Bredbenner, 2006), weight gain (Kelly, 2009; Jeffery, 2007) and bigger waistlines (Wansink, 2008; Schwartz and Byrd-Bredbenner, 2006) has been seen in observational studies involving both children and adults. 1.4 billion people, constituting 10% of global adult population in 2008 and 35 million children in 2010 were either obese or overweight globally (WHO, 2012b). Obesity, according to Lewington et al, (Boateng et al 2007) reduces life span of people with a body mass index (BMI) of 30 kg/m 2 by two to four years a nd in those with BMI of 40 kg/m 2 by eight to ten years.

### Obesity and the incidence of non-communicable diseases

Some risk factors for the development of NCDs are increased overweight and obesity due to unhealthy dietary and nutritional practices (Popkin, et al, 2012; WHO, 2012b; Cecchini, et al., 2010; Public Health Agency of Canada, 2010; Amoah, 2003). NCDs, defined as noncontagious, noninfectious chronic diseases that progress slowly over a long period such as cancer, cardiovascular diseases and diabetes mellitus and have beejL linked to unhealthy lifestyles such as poor dietary habits (WHO, 2011; Public Health Agency of Canada, 2010) are the highest contributors to global morbidity and mortality (WHO, 2005). NCDs have been linked to unhealthy lifestyles such as poor dietary habits (WHO, 2011); they are the leading cause of death with an estimated death of 36 million out of 57 million deaths in 2008 (WHO, 2010). The principal NCDs implicated in these deaths include cardiovascular diseases (17 million deaths, or 48% of NCD deaths) and diabetes (1.3 million deaths) (WHO, 2010). The Framingham Heart Study in the 1960s demonstrated a prospective relationship between body weight and hypertension (Lewis et al, 2013). The prevalence of cardiovascular disease (CVD) risk factors in West Africa is on the increase and obesity is one of them. A study conducted in Ghana and Nigeria reported a hypertension prevalence of 19.3 - 54 with overweight and obesity ranging between 20% to 62% and 4% to 49%, respectively. The study recorded only slight differences between rural, jirban, semi-urban, and mixed populations (Commodore Mensah et al, 2014). Nearly 55.2% of Ghanaian adults resident in Accra are either overweight or obese; about J3% of Ghanaians are overweight or obese (Ofori-Asenso et al, 2016). According to WHO, cardiovascular disease (CVD) is one of the top causes of death in Ghana and estimates that people aged between 30 to 70 years have a 20% risk of dying from CVD, cancer, diabetes and or of chronic respiratory disease (Boslaugh, 2011). Unfortunately, most people do not know what makes up a portion and this has been linked to their inability to control how much they eat (Byrd-Bredbenner & Schwartz, 2004). Portion size management of various food components have proved to be a very key behavioral element for weight management, (Penaforte et al., 2014) and that, cutting down portion sizes of particular foods at specific mealtimes has been shown to decrease day-to-day energy intakes (Lewis et al, 2015).

### Portion size in relation to undernutrition and incidence of NCDs

When smaller than recommended portion of food is consumed by children, pregnant women and women of child-bearing age, the result is undernutrition (Black et al., 2008; Benson and Shekar, 2006). A risk factor for intrauterine growth retardation (IUGR) is maternal undernutrition. Intrauterine growth retardation has been associated to low birth weight in infants, stunting and wasting, underweight and micronutrient deficiencies (Rehana et al, 2013), most of which are not reversible (Black et al, 2008). These infants and children are at risk of developing the metabolic syndrome, risk factors for NCDs (Barouki et al, 2012). According to Alberti et al, (2006), metabolic syndrome is defined by The International Diabetes Federation (IDF) as ‘a cluster of the most dangerous heart attack risk factors: diabetes and pre-diabetes, abdominal obesity, high cholesterol and high blood pressure‘. The WHO’s child growth programme recommends that when feeding the moderately wasted, stunted or underweight child from six months to twenty-four months, appropriate portion sizes is very important. This may ensure that adequate catch-up growth is realized during nutritional intervention strategies (Ashworth and Ferguson, 2008).

### Portion size control in reduction in incidence of NCDs

A reduction in consumption of larger portions of energy dense foods adds up over time and leads to an overall lower energy intake over the period (Rolls, Roe, et al, 2006). A decrease in the consumption of bigger portion sizes of foods lead to bigger losses of weight and fat with the corresponding reduction in the risk of cardiovascular disease (Hannum et al, 2004) and the risk of NCDs (Oguma et al, 2005).

### FoodEslNel

Problem Formulation

The Actual Food Estimation task is a Regression problem, where the input is a set of attributes on demographics, socioeconomic status, food type and perceived food amount and the output is an estimation of the actual amount of food. Our model’s objective during training is to optimise the Mean Absolute Error(MAE) loss

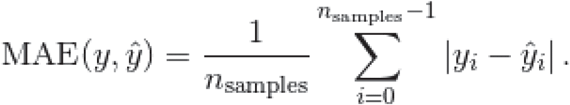

where the 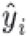 is the predicted output for *i*-th sample *y_i_*. is the (correct) target output computed over *n_samples_*

FoodEstNet uses XGBoost (Chen T. & Guestrin C, 2016), which is an implementation of Gradient Boosting (Friedman, 2001) that uses Hessian Matrix, a higher-order approximation, which helps it learn better tree structures, and it includes an extra randomization parameter, ie. column subsampling that helps it reduce correlation of each tree even further. Prior to choosing XGBoost, multiple Machine Learning algorithms were applied to the problem, including Linear Regression, Ridge, Lasso, Elastic Net, Bagging, Random Forest, Extra Trees, K Nearest Neighbors, Decision Tree, Gaussian Process, Support Vector Machine, Multilayer Perception(Neural Networks/Deep Learning) along with XGBoost. The XGBoost(with Early Stopping) model scored considerably higher than the other models. Lasso, Linear Regression, Ridge and Support Vector Machine also had decent high scores, the rest of the models had poor scores.

### Data

The dataset used for training and testing our model is from a cross-sectional study originally for the paper ‘Assessment Of Factors Affecting Perceived And Actual Carbohydrate Portion Sizes Among An Adult Population In Accra’ (Joseph B Danquah, 2017). The study is a cross-sectional study involving adults who are 18 years and above. A suburb was randomly selected from each of the four (4) income zones. Furthermore, a church and a mosque were randomly sampled rom each of the chosen suburbs. Convenient sampling was then used in selecting the participants from the churches and mosques for the study. Participants were then conveyed to a neutral ground for the study.

Participants’ perception of portion sizes served of some commonly consumed carbohydrate foods and actual portions sizes served were obtained and analyzed in relation to gender, age and BMI. Participants’ socio-economic and demographic data were also collected and analyzed. Four hundred (400) adults were assessed on carbohydrate portion size perception and actual portion sizes they consume. Twelve commonly consumed carbohydrate foods made up of 50% grain and cereal group, 33.3% roots, tubers and plantain group and 16.7% of beverage and sugar group were collected. The carbohydrate foods that were collected are chocolate drink, white bread, boiled rice, Ga kenkey, granulated sugar, corn porridge, waakye, gari, boiled yam,boiled green plantain, banku and fufu. There were 212 (53%) males and 188 (47%) females.

### Estimation of portion sizes of carbohydrate food

Handy measures were chosen for the study and: these were handy measures identified by Owusu et al, (1995) as commonly used in the Ghanaian household and used in the dietetic clinics to assess portion sizes. The handy measures used comprised of ladles, stewing spoons, tablespoons, dessert spoons, teaspoons, sardine tin, small tomato puree tin and 450 g margarine tin. Participants were allowed to select handy measures from the handy measures presented, based on the type they usually use to serve the selected carbohydrate foods. Twelve commonly consumed carbohydrate foods were presented in the questionnaire. Participants wrote or were helped to write on a questionnaire (dietary perception survey form), against each carbohydrate food, in handy measures, how much of each carbohydrate food they believed was their portion sizes. They were then presented with the selected carbohydrate foods and asked to serve their perceived portion sizes of each carbohydrate food using the same handy measure tool. A comparison of perceived portion size and actual portion size of each carbohydrate food was made.

### Pre-Testing Questionnaire

The socio-demographic questionnaire was tested among 20 adults of which 10 were females and 10 were males. These adults were conveniently selected to meet the study criteria from the University of Ghana, Legon campus. The questionnaire requested from the participants their socio-demographic information such as age, gender, ethnicity, religion, marital status, education level, health status, assessment of food aid; anthropometric measurements; height, weight and BMI.

### Socio-economic and demographic data

Data on socio-demographic variables such as gender, educational status, age, marital status, family size, occupation, average income, religion and ethnicity were collected using semi-structured questionnaire. Anthropometric assessment In addition to nutritional and dietary intake assessments, and assessment of laboratory data, anthropometric assessment is very much needed in studying the relationship among diet, nutritional status and health (http://www.cdc.gov/growthcharts). According to the National Health and Nutrition Examination Survey (NHANES), assessment of nutritional status requires a series of stature, weight, and other anthropometric dimensions. Researchers in various clinical disciplines such as nutrition and dietetics, cardiovascular health, physiotherapy and occupationartioal health make use of anthropometric data to make a diagnosis and examine health status trends (Barbara J Rolls et al, 2002).

The anthropometric measurements taken for the purpose of this research included weight in kilogram, (kg) and height in centimeters,(cm). The body mass index (BMI) of each participant was then calculated as weight in kilograms, divided by height in meters squared (kg/m2). Prior to calculation of BMI, unit measurement of height in centimeters was converted to meters. The anthropometric data obtained were compared to reference standards of an adult as defined by the World Health Organisation, WHO (2016). Weights and heights were recorded by researcher and trained research assistants for all participants.

### Weight

The weights of participants were measured by the researcher and trained research assistants at the time of questionnaire distribution in duplicate to the nearest 0.1 kg, using a Floor Omron BF508 Body Composition Monitor, with subjects standing upright without shoes. Participants were in light clothing, and were kindly asked to remove jackets, shoes, belts, wallets, keys and other objects including mobile phones before standing on the scale. The BMI was calculated for each participant using his or her height and weight data collected.

### Height

The heights of participants were measured also at the time of questionnaire distribution in duplicate to the nearest 0.1 cm with a Seca 213 portable stadiometer. Participants stood upright against the stadiometer with the back of their heels and occiput touching the wall of height measure.

### Collection of commonly consumed carbohydrate foods

The commonly consumed carbohydrate foods collected were composed of foods from cereals and grains group, roots, tubers and plantain groups and then, sugar and beverage group. Commonly consumed carbohydrate foods were defined by Boateng (2014) as carbohydrate based foods that were consumed at least two times for two week-days and one week-end when meals were recorded. These carbohydrate foods informed the choice of foods used in this study. Any carbohydrate based food that did not fit this criterion was not included in the food items collected. The carbohydrate foods that were collected are chocolate drink, white bread, boiled rice, Ga kenkey, granulated sugar, corn porridge, waakye, gari, boiled yam, boiled green plantain, banku and fufu.

### Data Preparation

The collected data was then prepared and changed to best suit Machine Learning. The problem was rephrased to estimate how much of a specific food type does a person consume, given the food type, perceived consumed amount of that food type and all other attributes as input. To accomplish this, we merged all 12 seperate food types attributes into one attribute, Food Type, by transposing the columns of data samples. The results of this was a 4,800 data samples out of the original 400 data samples. The benefit of this is it gave us a bigger dataset which is very important to developing Machine Learning models. And also, instead of having our final trained model require all 12 food types as input from the user before outputting a sequence of estimations or even having to train 12 separate models for each food type with just 400 samples each, we can rather have a more universal model capable of estimating any of the 12 main food types. We also removed Selected Handy Measure column because it doesn‘t serve any purpose in our model, it doesn’t have any influence on the estimates.

### Data Analysis

We ended up with a dataset that had 4,800 data samples and 12 columns. The 12 columns were; Sex, Age, Ethnicity, Marital Status, Education, Average Monthly Income, Mental & Physical Health, BMI, Religion, Food Type, Perceived Amount, Actual Amount. The table below shows the values for each of the columns and the Numerical Encodings used for our modelling;

The table below shows the values for each of the columns and the Numerical Encodings used for our modelling;

| Attributes | Values | Encoding |
| --- | --- | --- |
| Sex | Male | 1 |
|  | Female | 2 |
| Age | Continuous | Continuous |
| Ethnicity | Akan | 1 |
|  | Ga/Dangme | 2 |
|  | Ewe | 3 |
|  | Guan | 4 |
|  | Mole /Dagbani | 5 |
|  | Hausa | 6 |
|  | Other | 7 |
| Marital Status | Never married | 1 |
|  | Currently married or cohabiting | 2 |
|  | Separated/divorced | 3 |
|  | Widowed | 4 |
| Education | Middle/junior high school | 1 |
|  | Secondary /Senior secondary school | 2 |
|  | Post-secondary/tertiary level completed | 3 |
|  | Post-graduate degree | 4 |
| Average Monthly Income(GH Cedis) | < 200.00 | 1 |
|  | 200.00 to < 500.00 | 2 |
|  | 500 to < 1000 | 3 |
|  | 1000 to < 1500 | 4 |
|  | ≥ 1500 | 5 |
| Mental & Physical Health | Excellent /very good | 1 |
|  | Good | 2 |
|  | Fair/moderate | 3 |
|  | Poor | 4 |
| BMI | Continuous | Continuous |
| Religion | Christian | 1 |
|  | Moslem | 2 |
|  | Traditionalist | 3 |
|  | Other | 4 |
| Food Type | Yam | 1 |
|  | Waakye | 2 |
|  | Sugar | 3 |
|  | Rice | 4 |
|  | Plantain | 5 |
|  | Kenkey<br>Gari<br>Fufu<br>Porridge<br>Chocolate<br>Banku<br>Bread | 6<br>7<br>8<br>9<br>10<br>11<br>12 |
| **Perceived Amount** | Continuous | Continuous |
| **Actual Amount** | Continuous | Continuous |

And the table below shows some general Analysis of the attributes

|  | Mean | Standard Deviation | Min | 25% | 50% | 75% | Max |
| --- | --- | --- | --- | --- | --- | --- | --- |
| **Sex** | 1.4850 | 0.552122 | 1 | 1 | 1 | 2 | 6 |
| **Age** | 36.72250 | 13.381995 | 18 | 27 | 35 | 43 | 81 |
| **Ethnicity** | 2.42750 | 1.75795 | 1 | 1 | 2 | 4 | 7 |
| **Marital Status** | 1.80250 | 0.763947 | 1 | 1 | 2 | 2 | 4 |
| **Education** | 2.22250 | 0.963425 | 1 | 1 | 2 | 3 | 4 |
| **Average Monthly Income** | 2.43250 | 1.095762 | 1 | 2 | 2 | 3 | 5 |
| **Mental & Physical Health** | 3.61750 | 0.520817 | 1 | 3 | 4 | 4 | 4 |
| **BMI** | 25.803750 | 5.848503 | 12 | 21 | 25.3 | 29.63 | 42.8 |

### Model Architecture and Training

The dataset was splitted into Training set and Test set, with a 9:1 ratio (4320, 480), respectively. The split was done stratified with the Food Type so each Food Type still gets equally split with a 9:1 ratio between the Training and Testing sets. This is to prevent under training on some food types compared to others. Both input values of the Training and Testing attributes were preprocessed with Standardization which centers the data by removing the mean value of each feature, then scales it by dividing non-constant features by their standard deviation. The XGBoost model was then trained with the Training set, with an Early Stopping set to 52 rounds in order to manage computation and prevent overfitting. The Mean Absolute Error(MAE) function was used used as evaluation metric during training.

Our XGBoost model was developed with the standard parameters, booster: gbtree, eta(learning_rate) = 0.3, gamma(min_split_loss) = 0, max_depth = 6, min_child_weight = 1, max_delta_step = 0, subsample = 1, colsample_bytree = 1, colsample_bylevel = l, lambda(reg_lambda) = 1, alpha(reg_alpha) = 0, sketch_eps = 0.03, scale_pos_weight = 1, refreshjeaf = 1, processjype = ‘default’, grow_policy = ‘depthwise’, objective = regdinear, base_score = 0.5.

### Model Testing and Results

The model was then tested with the Test set, which contained 480 samples. Explained Variance Accuracy score and Root Mean Square Error(RMSE or RMSD) metrics were used to measure the performance of the model. With their respective formulae being:

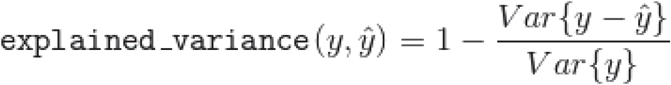

where 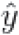 is the estimated target output, *y* the corresponding (correct) target output, and *Var* is Variance, the square of the standard deviation. And,

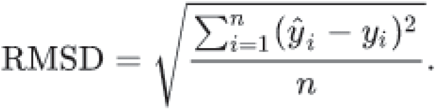

where the 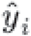 is the predicted output for observations *i* and *y_i_* is the (correct) (correct) target output computed for *n* different predictions

### Conclusion And Outlook

We developed the model FoodEstNet that estimates True Food Consumption by training and testing the model with a dataset that had 4,800 data samples and 12 columns, one being the output column. We‘ve shown it is possible to use Machine Learning/Artificial Intelligence to estimate the true amount of food consumed by people. With a bigger and broader study for larger datasets and additional research, we’re confident models could be develop with high accuracy scores and performances that they could be used in the real world by Nutritionists, Dieticians and people in general alike to accurately estimate Food Consumption for foo portion size control and nutritional management of diseases. We’d also like to in the near future explore developing models that can estimate calorie amounts in mainly Ghanaian dishes using only images of such dishes.

### Limitations

Limitations come by, as a result of specifics of the experimental technique. Some confoundingfactors may have had an effect on participants’ abilities to correctly estimate portion sizes. However, they could not be effectively controlled for in this study and thus may influence the proportions of the different degrees of estimation reported, hese factors are outlined below: Actual portion size estimation was done in the open with limited insurance of privacy. Therefore, some participants might either increase or decrease their portions to avoid intimidation. A participant’s level of like and dislike for certain carbohydrate foods may affect the portion size estimations. Confounding factors such as temperature and heat loss in the form of vapour might have affected the size and weight of some carbohydrate foods, hence their estimations. Also, the limited amount of samples, 480 per Food Type, isn’t quite sufficient for effectively training Machine Learning model, a bigger dataset is likely to yield better performance and score.

## Acknowledgements

We’d like to acknowledge all the participants of the cross-sectional study that gave us the dataset needed to embark on this research.

## Abbreviations

Al: Artificial Intelligence
NCD: non-communicable diseases.

